# Soil trace gas oxidizers divergently respond to short- and long-term warming

**DOI:** 10.1101/2025.11.18.689010

**Authors:** Thanh Nguyen-Dinh, Vojtěch Tláskal, Pok Man Leung, Magdalena Wutkowska, Justus Amuche Nweze, Katri Ylä-Soininmäki, Johanna Kerttula, Mathilde Borg Dahl, Andrea Söllinger, Tilman Schmider, Lukas Kohl, Christina Biasi, Andreas Richter, Tim Urich, Alexander T. Tveit, Chris Greening, Bjarni D. Sigurdsson, Anne Daebeler

## Abstract

The upland soil microbiome is dominated by aerobic bacteria that oxidize atmospheric trace gases, including CO, H_2_, and CH_4_. As a result, soils are the largest biological sink for these climate-active gases. Whether global warming will enhance or suppress these processes remains unclear. Here, we studied the warming responses of soil trace gas oxidizers by profiling natural geothermal gradients in a subarctic grassland with over 60 years of field warming at +6°C. We integrate field flux measurements, *ex situ* biogeochemical assays, metagenomics, and metatranscriptomics to determine ecosystem and cellular-level responses. Our results show that the oxidation of atmospheric CO and H_2_, but not CH_4_, increased with long-term warming due to higher cell numbers. However, at the cellular level, trace gas oxidizers, especially methanotrophs, tended to reduce gas consumption and transcription of gas-metabolizing enzymes in response to long-term warming. Our findings suggest that soils may remain a robust sink for trace gases despite lower per-cell activity. This work establishes a framework for interpreting the relationships between temperature and microbial trace gas oxidation on timescales relevant to Earth’s climate system.

## Introduction

Carbon monoxide (CO), hydrogen (H_2_), and methane (CH_4_) are the most abundant climate-forcing reduced gases in Earth’s atmosphere. CO and H_2_ are indirect greenhouse gases owing to their reactivity with atmospheric hydroxyl radicals; they have a 100-year global warming potential equivalent to 2 and 12 times that of carbon dioxide (CO_2_), respectively (1, 2). CH_4_ is the second most abundant greenhouse gas and has a 30-fold stronger warming potential than CO2 on a 100-year timescale (3).

Anthropogenic activities have increased emissions of all three gases since industrialization, accelerating global warming (4–6). Soil microorganisms play a key role in regulating the atmospheric budgets of these gases, acting as both sources and sinks. They constitute the single most important biological sink through microbial oxidation, contributing with 70–80%, 10–20%, and 3–5% of the annual atmospheric removal of H _2_ (7–9), CO (10, 11), and CH_4_ (12–14), respectively. While this microbial sink has helped to keep CO and H_2_ levels in the atmosphere relatively constant [mixing ratios of 0.09 ppmv for CO (11) and 0.53 ppmv for H_2_ (9)], the relatively weak sink for CH_4_ has not effectively counterbalanced emissions and atmospheric levels have increased almost threefold (to 1.91 ppmv in 2021) compared to preindustrial concentrations (5). The microbial trace gas sink, therefore, provides a significant, albeit not complete, mitigation of the climate impact of these gases.

Trace gas oxidizers are taxonomically diverse and abundant inhabitants of different soils across the globe, with an average of 56, 39, and 1% of the total soil microbial community predicted to use atmospheric CO, H_2_, and CH_4_ as energy sources, respectively (15, 16). They utilize specialized enzymes, specifically form I [MoCu]-CO dehydrogenases (17), high-affinity [NiFe]-hydrogenases (18), and particulate CH_4_ monooxygenases (19), to extract energy from the ubiquitous yet diluted atmospheric substrates. Atmospheric CO and H_2_ primarily provide microbes an important continuous energy supply to persist under energy limitation (20–22); in-depth studies have revealed that microbes upregulate expression of high-affinity hydrogenases and CO dehydrogenases in direct response to organic carbon deprivation, and that these enzymes greatly enhance survival during starvation (23–26). Most trace gas oxidizers use trace gases at atmospheric concentrations to support mixotrophic growth with organic or alternative energy sources (22, 27, 28). Remarkably, some oligotrophic bacteria are even able to grow on atmospheric energy sources alone, as recently demonstrated for several atmospheric methanotrophs (27, 29). Together, their activities are responsible for a net annual uptake of 250 Tg of CO, 60 Tg of H_2_, and 31 Tg of CH_4_ from the atmosphere [(30) and references therein; (14) for CH_4_].

Despite the abundant and active nature of soil microbes in regulating atmospheric trace gas levels, the impact of climate change on this important soil ecosystem service is unknown. Global warming is one of the largest threats to crucial soil functions mediated by microorganisms, including organic carbon storage (31, 32). Previous studies have shown significant and persistent soil organic carbon (SOC) depletion after warming of ≤6°C (33–36). This SOC loss is driven by warming-induced acceleration of microbial activity, but is stabilised by a decline in microbial biomass in the long term as the microbial biomass equilibrates with the depleted SOC stores (33, 35, 37). The new long-term steady state of topsoil in a warmer world is therefore predicted to be characterized by reduced microbial biomass, species richness, and SOC stocks (35). The shifts in microbial community structure of SOC decomposers and in resource availability may also influence the abundance and activity of trace gas oxidizers, yet this link remains poorly understood. Very little research is available to predict if the soil sink for climate-active trace gases will be maintained, decline, or even increase in a future climate (38, 39). Any change in this regard is likely to have far-reaching climatic, ecological, and biogeochemical ramifications, as outlined above. This is especially crucial as emission of climate-related gases are projected to increase, for example due to the transition to a hydrogen economy (fugitive H_2_ emissions) (40), intensifying wildfires (CO release from incomplete combustion) (41), positive feedback dynamics from wetlands (42), and growing agricultural demand (CH_4_ emissions from ruminants and rice paddies) (43). Moreover, a depletion of the chemical sink for trace gases (hydroxyl radicals) has been inferred to cause increased levels of atmospheric CH_4_ (44).

Here, we leveraged an *in situ*, natural warming study site (> 60 years) (45) to determine the response of soil CO, H_2_, and CH_4_ oxidizers to long-term warming. The study site harnesses the natural geothermal heat from the Mid-Atlantic Ridge, which is transmitted to the bedrock beneath a subarctic grassland, creating temperature gradients ranging from unwarmed to +6°C and above. This represents an unparalleled research site to assess long-term microbial responses of distinct functional groups and of the entire ecosystem in a region that holds substantial carbon stocks and is vulnerable to rapid temperature change (46). In this study, we systematically assessed the response of trace CO, H_2_, and CH_4_ oxidizers to both short-term (one month) and long-term (decades) soil warming. We quantified the fluxes of climate-active atmospheric gases in the field and used metagenomics and metatranscriptomics to characterize the present and active trace gas oxidizers in unwarmed and long-term warmed soil. We then used the same soil for a laboratory experiment to quantify the potential soil uptake rates of trace levels of CO, H_2_, and CH_4_ in unwarmed, short-term warmed (with only days of warming in the lab), and long-term warmed soil (with decades of warming *in situ*). We hypothesized that i) trace gas oxidizers will show increased oxidation rates in both the short- and the long-term warming condition due to accelerated activity and as a response to decreased SOC availability, respectively; ii) trace gas oxidizers will increase in relative abundance in the long-term warmed soil, as they are able to sustain themselves better under SOC depletion caused by warming than microbes not able to utilize trace gases; and iii) at the ecosystem level, warmed soil will show an increased sink capacity for CO, H_2_, and CH4 in both the short- and the long-term.

## Results

### Long-term warming enhances the CO and H_2_ but not the CH_4_ sink

We profiled four replicated temperature transects, comparing unwarmed and long-term warmed soil (6°C above ambient, Fig. S1) for differences in key microbial (cell count, microbial carbon, microbial nitrogen) and physico-chemical (pH, moisture, dissolved organic carbon, total dissolved nitrogen) parameters (Fig. S2, Table S1). The number of bacteria and archaea, as determined by 16S rRNA qPCR quantification integrated with metagenome data, was statistically similar between the unwarmed and warmed soils; however, three out of the four investigated warming transects showed an increase (Fig. S2). Additionally, the warmed soil exhibited a higher respiration activity, with *in situ* soil CO_2_ emission (751.7 ± 123.2 mg CO_2_ m⁻^2^ h⁻¹) at a significant 1.6-fold faster rate than from unwarmed soil (460.7 ± 9.8 mg CO_2_ m⁻^2^ h⁻¹) (Fig. 1A, Table S2). However, soil CO_2_ concentrations along the soil depth profile were comparable in the long-term warmed and unwarmed soil with significantly higher concentrations in the deepest analysed layer of 20 cm (Fig. 1C). The corresponding δ¹³C-COC-CO_2_ values became progressively lighter with increasing depth with the exception of two warmed plots at which we observed a slight increase in δ¹³C-COC-CO_2_ values with depth suggesting a slight inflow of geogenic CO_2_ (Fig. 1E, Table S3).

**Figure 1.**
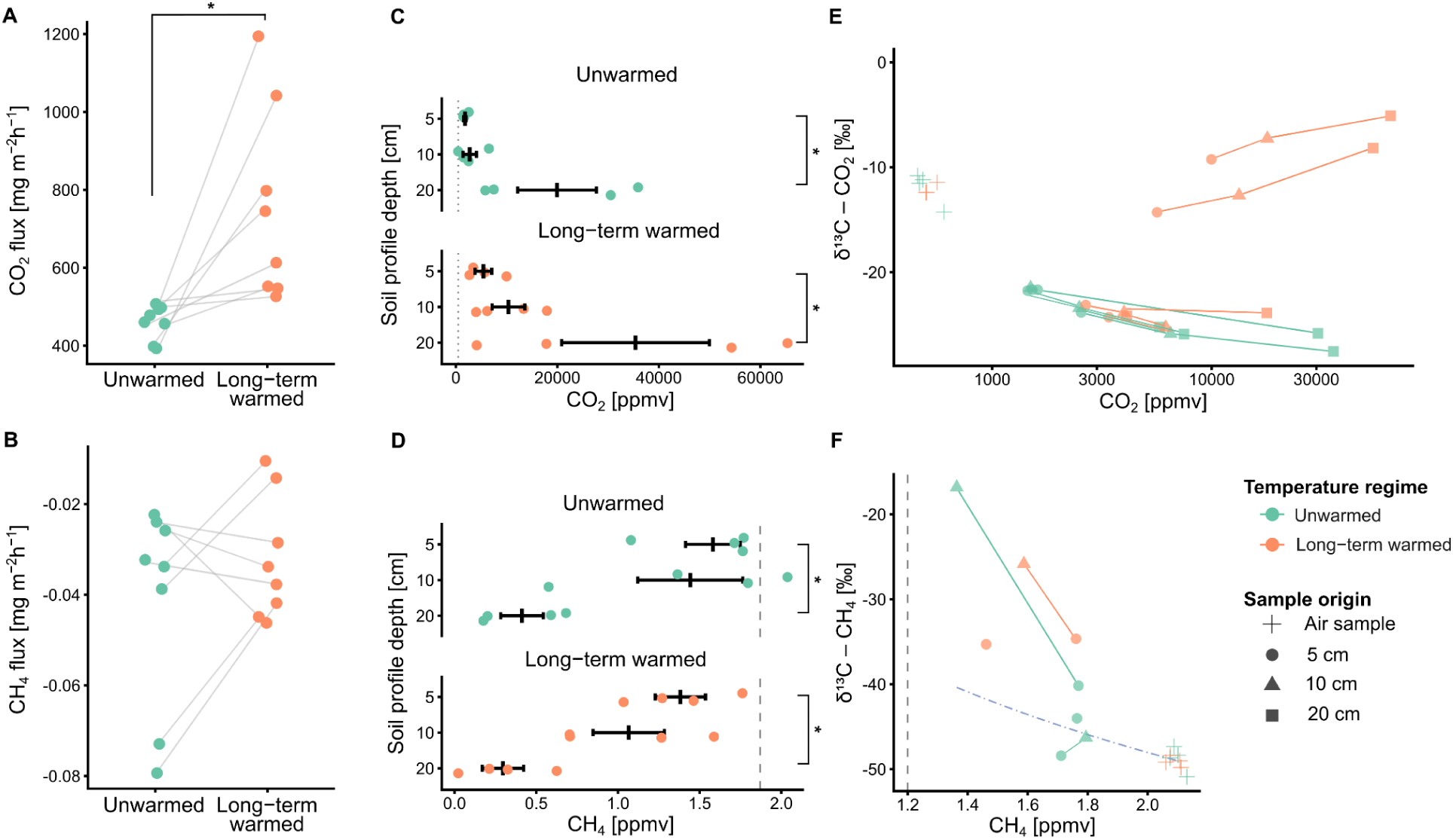
Soil-atmosphere gas fluxes, soil gas concentrations, and C isotope signatures. *In situ* flux of CO_2_ (**A**) and CH4 (**B**) from unwarmed and long-term warmed soil. Individual data points from two consecutive days are shown, with lines connecting pairs within a single warming transect used for statistical testing. Concentrations of CO _2_ (**C**) and CH_4_ (**D**) in unwarmed and long-term warmed soil at different depths (5, 10, and 20 cm, mean ± SE are depicted by black lines). Vertical dotted lines denote the average atmospheric CO2 and CH4 concentration measured on site. Relationship between CO_2_ (**E**) and CH_4_ (**F**) concentrations and C isotope values in unwarmed and long-term warmed soil at different depths. The dot-dashed line in (**F**) represents a model following a Rayleigh distillation function with a fractionation factor (alpha) of 1.020. The vertical dashed line in (**F**) denotes the detection limit for δ^13^C-CH_4_ in samples with <1.2 ppmv CH_4_. Significant differences and trends are depicted by square brackets and an asterisk or dot, respectively.

In contrast to the communities mediating organic carbon respiration, soil methanotrophic activity was not stimulated by long-term warming. There was an overall 22% decrease in *in situ* atmospheric net CH_4_ uptake in warmed (0.032 ± 0.007 mg CH_4_ m⁻^2^ h⁻¹) compared to unwarmed plots (0.041 ± 0.012 mg CH_4_ m⁻^2^ h⁻¹), though rates varied considerably between plots (Fig. 1B, Table S2). CH_4_ concentrations decreased similarly with depth in the soil pore space of both unwarmed and warmed soil, and δ¹³C-COC-CH_4_ signatures became increasingly heavier with soil depth (Fig. 1D, F, Table S3). A model estimate of δ¹³C-COC-CH_4_ values along the soil profile based on methanotrophic δ¹³C-COC-CH_4_ fractionation agreed well with the obtained data, indicating that CH_4_ oxidation can be assumed as the reason for the observed decrease in CH_4_ in deeper soil layers (Fig. 1F). These patterns suggest no biotic or geogenic CH_4_ sources were present below the study plots. Therefore, the measured net CH _4_ uptake likely corresponds to the gross CH_4_ oxidation activity of atmospheric CH_4_ oxidizers in the soil. This is further corroborated by the negligible abundance and transcription of key methanogenesis genes in the obtained metagenomes and metatranscriptomes (Fig. S3, Table S4–S7). Logistical challenges prevented an equivalent quantification of soil CO and H_2_ fluxes in the field; however, we were able to detect *in situ* uptake of both gases by unwarmed and warmed soil (Table S8). As a support to this unreplicated observation, a diffusion model for the studied trace gases was built, showing the possible uptake of CO and H_2_ by the upper soil layer and likely depletion of these gases within the top 10 cm, while CH_4_ was predicted to diffuse deeper and only be depleted at a depth of >20 cm (Table S9). This aligns well with our measured CH _4_, H_2_, and CO uptake data (Tables S2, S8) and provides confidence in the estimated gas flux across the soil profile.

Soil microcosms were prepared to measure the *ex situ* oxidation of trace-level CO, H_2_, and CH_4_, with soil from unwarmed and long-term warmed plots incubated at their respective average summer temperature of 13°C and 19°C, respectively. A short-term warming treatment (incubation at 19°C for 40 days) with soil from unwarmed plots was also included to evaluate the effect of a warming pulse. Both atmospheric H _2_ and CO oxidation rates were significantly faster in long-term warmed soil (0.72 and 0.076 nmol h^-1^ gDW^-1^, respectively) than in unwarmed soil (0.25 and 0.034 nmol h^-1^ gDW^-1^, respectively) (Fig. 2A–B, Table S10–S12). In contrast, CH_4_ oxidation rates in the microcosms remained similar (Fig. 2C), in line with the unchanged *in situ* soil CH_4_ uptake of unwarmed and warmed soil (Fig. 1A). These data imply increased soil sink capacities for CO and H_2_, but not for CH_4_ uptake in the long term. Short-term warming of soil, on the other hand, did not lead to changes in the consumption rates of any of the three gases (Fig. 2A–C, Table S11–S12), suggesting that the size of the soil trace gas sink is resilient to short-term warming.

**Figure 2.**
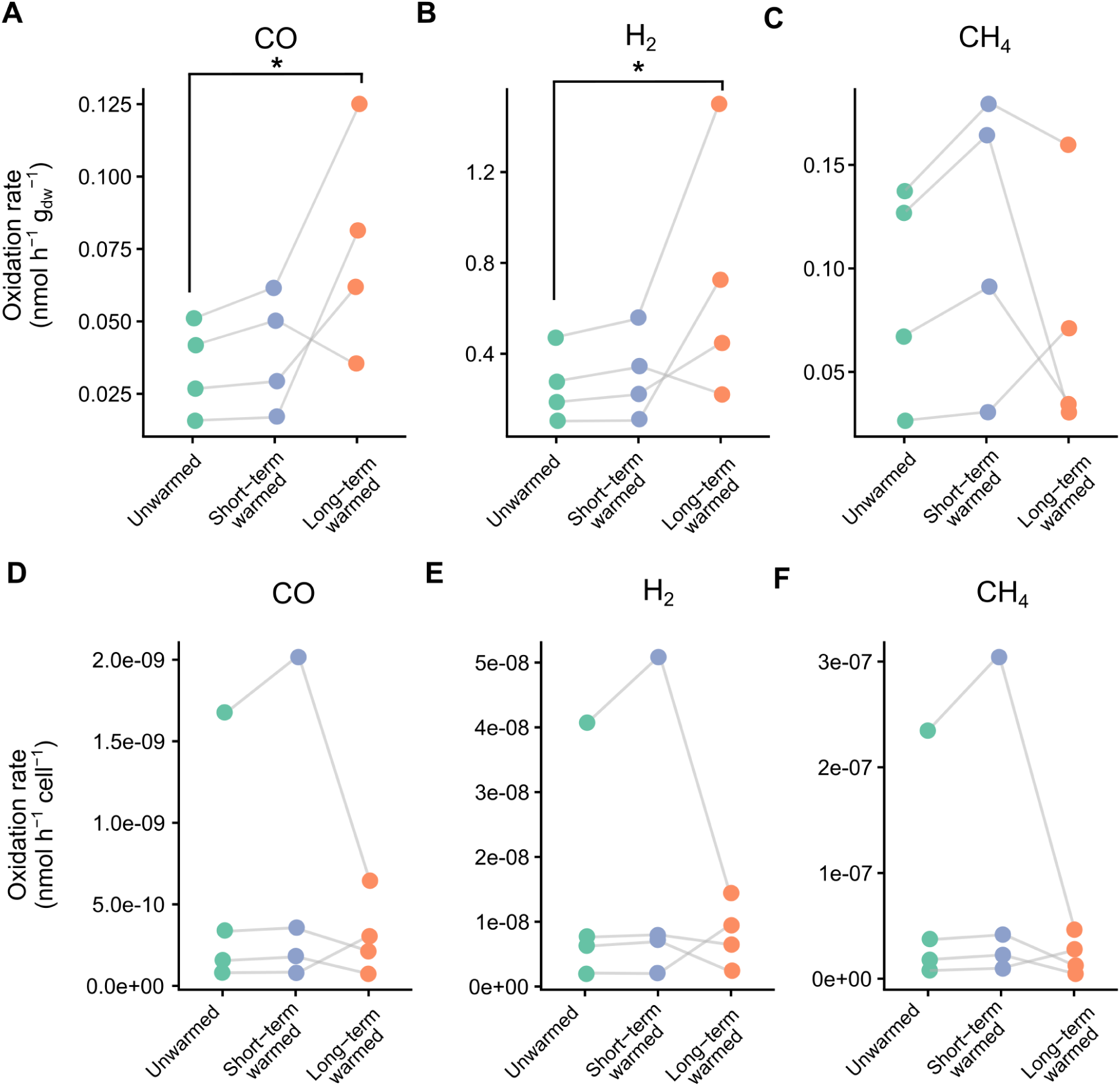
*Ex situ* trace gas oxidation rates. Bulk soil oxidation rates (**A, B, C**) and cell-specific oxidation rates (**D, E, F**) of CO, H_2_, and CH_4_ obtained from unwarmed, short-term warmed, and long-term warmed incubation treatments. Individual data points are shown together with lines connecting samples from the same warming transect. DW stands for dry weight; asterisks denote significant differences between treatments.

### Warming impacts the relative abundance and community composition of trace gas oxidizers

To investigate how long-term warming modulates the populations of trace gas oxidizers, we obtained metagenomes from samples of both unwarmed and long-term warmed plots. Additionally, we included data from previously obtained metagenomes of the same plots into the analyses. A gene-centric approach was applied to profile the community abundance of 51 metabolic marker genes, specifically form I CO dehydrogenases, hydrogenases, and CH_4_ monooxygenases, which catalyze the oxidation of CO, H_2_, and CH_4_, respectively (Fig. 3, Fig. S3, Table S4–S5). Consistent with our prior survey on temperate soils (*15*), hydrogenases (primarily group 1h [NiFe]-hydrogenases) and [MoCu]-CO dehydrogenases (*coxL*) that mediate atmospheric uptake during long-term survival were encoded by most microbial community members, whereas CH_4_ monooxygenases (*pmoA*, *mmoX*) were relatively rare, under both temperature conditions (Fig. S3). Although not significant, warmed soil harboured a higher abundance of [NiFe]-hydrogenases (warmed: 0.69, unwarmed: 0.56 copies/cell, based on the abundance of hydrogenase genes related to the summed count of 14 single-copy marker genes), whereas CH_4_ monooxygenases decreased by an average of 32% (warmed: 0.012, unwarmed: 0.017 copies/cell) (**F**ig. 3A). No notable relative abundance shifts in the CO dehydrogenases (**F**ig. 3A) or any of the other profiled metabolic marker genes were observed (Fig. S3, Table S5).

**Figure 3.**
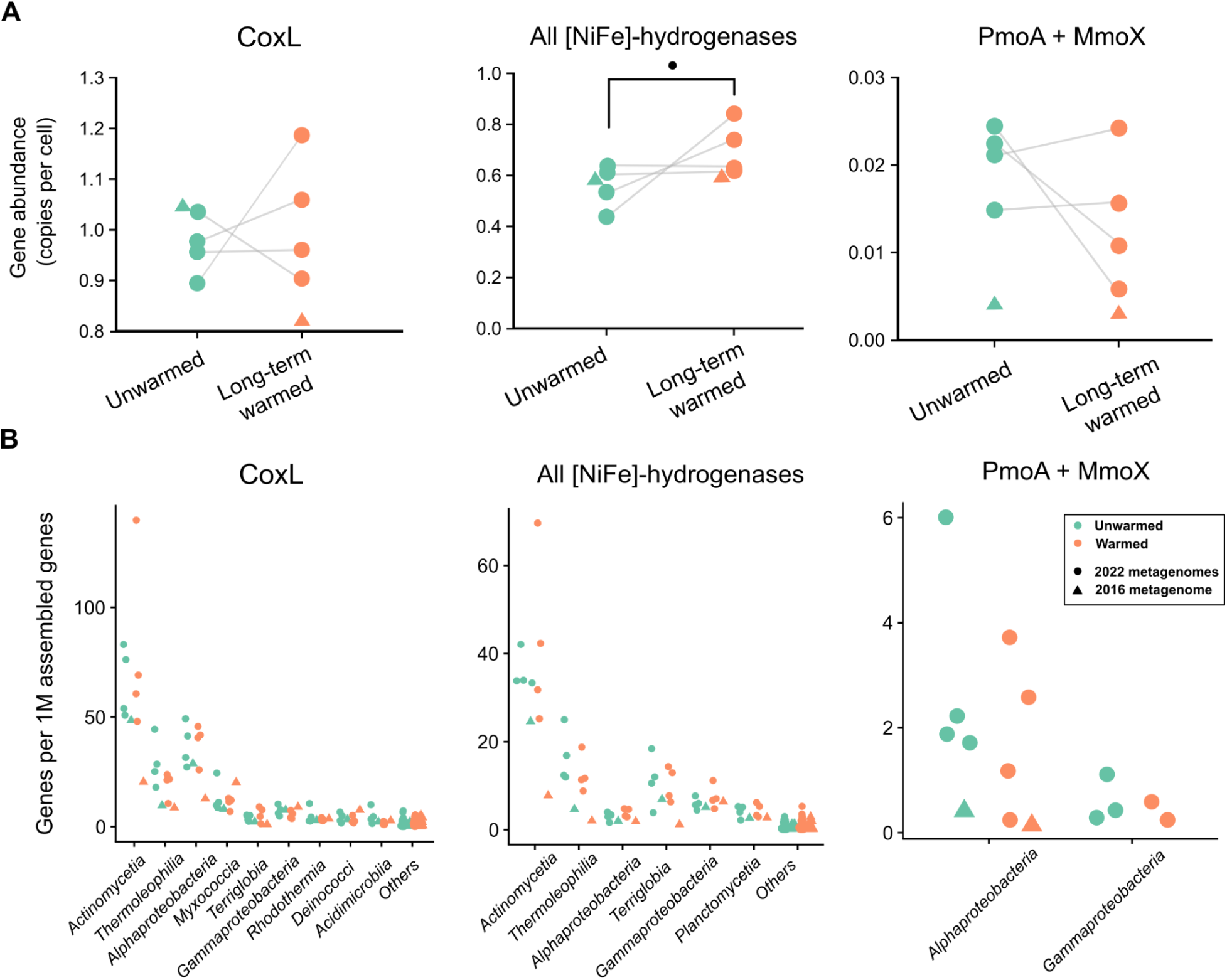
Relative abundance shifts of trace gas oxidizing gene composition caused by warming. (**A**) Relative abundance of marker genes, calculated as gene copies per cell. (**B**) Relative abundance values of marker genes in relative proportion to total assembled gene counts for the most abundant taxa are displayed. Low-abundant taxa are grouped as “Other” for *coxL* (overall abundance average <3.2% per taxon) and [NiFe]-hydrogenases (<4.3%). Data shown are from this study (circles; n= 4) and from public metagenomic short reads (BioProject ID PRJNA746424, triangles; n= 1). Lines indicate pairs within a single warming transect that were used for statistical testing. Significant trends are depicted by square brackets and a dot.

We additionally analysed shifts in the relative abundance of major soil prokaryotic taxa mediating trace gas oxidation by profiling the taxonomic affiliation of the trace gas oxidation marker genes from the metagenomic assembly. Based on this analysis, the richness of CO (180 ± 27 unique best hit genomes), H_2_ (160 ± 36), and CH_4_ (4.0 ± 0.8) oxidizer communities were not affected by warming. [NiFe]-hydrogenases and CO dehydrogenases were widespread, encoded by 22 and 13 phyla, respectively, and predominantly associated with the bacterial classes *Actinomycetia*, *Thermoleophilia*, and *Alphaproteobacteria* (Fig. 3B, Table S13). Actinomycetal CO and H_2_ oxidizers showed a trend of increased relative abundance with long-term soil warming, while Thermoleophilia groups overall decreased. CH_4_ monooxygenases were mainly assigned to alphaproteobacterial (primarily *Methylocella* spp.; *Methylocapsa* spp. in the recent GTDB v226) and to a lesser extent to gammaproteobacterial taxa, and both groups showed a trend of decreased relative abundance with warming (Fig. 3B). This observation is consistent with metagenomic community profiling based on 16S rRNA genes by phyloFlash (Table S14).

To gain deeper insights into the mediators of trace gas oxidation, we performed metagenomic assembly and binning, yielding 250 high- and medium-quality metagenome-assembled genomes (MAGs; dereplicated at 95% ANI) (Table S15). Of these, 110 bacterial MAGs spanning 12 phyla held the genomic potential for trace gas oxidation, namely 88, 45, and 1 MAGs encoding the enzymes for atmospheric CO, H_2_, and CH_4_ oxidation, respectively (Fig. 4, Table S16). To determine whether long-term warming affected the abundance of distinct trace gas oxidizers, we mapped the MAGs to the metagenomic short reads from the unwarmed and warmed soils (Table S17). In line with the results of the short-read and assembled-read analyses outlined above, more than half of the CO- and H_2_-oxidizing MAGs were relatively enriched in the warmed soil (50 and 27 MAGs, respectively), while the sole CH_4_-oxidizing MAG (*Methylocella* spp.) decreased by 4.3-fold (Fig. 4). However, few of these differences were statistically significant (after correcting for multiple comparisons) given the considerable variability in community composition between the plots. Finally, there was an increase of summed relative abundance of H_2_-oxidising MAGs (1.3 ± 0.2% unwarmed, 2.0 ± 0.1% warmed) in warmed soil, whereas the summed relative abundance of all MAGs was unchanged (8.7 ± 2.0% unwarmed, 11.3 ± 0.6% warmed, Fig. S4). Altogether, these analyses suggest diverse CO and H_2_ oxidizers increased in relative abundance with warming, whereas methanotrophs decreased, which aligns well with the *in situ* and *ex situ* activity data.

**Figure 4.**
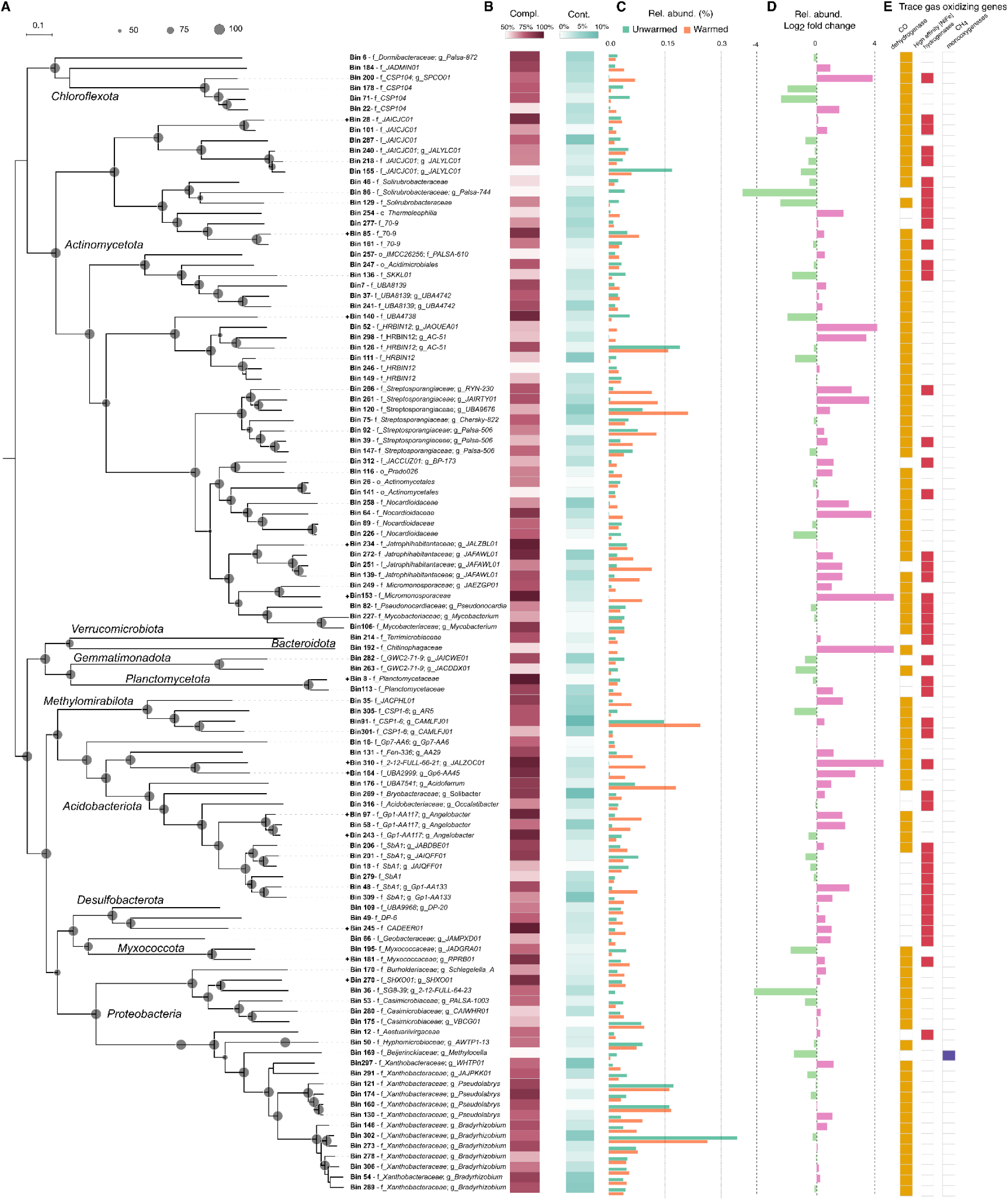
Phylogenomics and differential abundance of non-redundant trace gas oxidizer genomes. **(A)** The phylogenomic tree of 120 medium- and high-quality metagenome-assembled genomes (MAGs) was constructed using approximately maximum likelihood estimation with the LG+F+I+G4 model. The scale bar represents 10% sequence divergence, and grey circles at nodes indicate bootstrap support. High-quality MAGs (completeness > 90%, contamination < 5%) are labeled with stars. All other MAGs have medium quality (completeness > 50%, contamination < 10%). **(B)** Completeness and contamination values of MAGs. **(C)** Average relative abundance (%) of trace gas oxidizing MAGs in unwarmed and warmed soils. **(D)** Log_2_ fold change in relative abundance between warmed and unwarmed soil (positive values = enrichment in warmed soil). **(E)** Presence/absence of genes involved in trace gas oxidation in MAGs. Type I CO dehydrogenase (CoxL) for CO oxidation (yellow), high-affinity [NiFe]-hydrogenases (group 1f, 1l, 1h and 2a) (red), and CH_4_ monooxygenases (PmoA+MmoX) for CH_4_ oxidation (blue).

### Cell-specific gene transcription and activity of trace gas oxidizers decrease with warming

We obtained the metatranscriptomes of the same soil samples to further determine the relative transcriptional activities of soil trace gas oxidizers (Fig. 5). As done for the metagenome analyses, we also included previously obtained metatranscriptomes from the same plots. For H_2_ oxidation, [NiFe]-hydrogenases were abundantly transcribed at both temperature conditions (Fig. S3, Table S18), but transcription ratios (average RNA:DNA ratios) were significantly lower in warmed soil, as compared to unwarmed soil (0.22 and 0.39, respectively) (Fig. 5B, Table S19–S20). For CO and CH_4_ oxidation, transcripts and transcription ratios for CO dehydrogenases and CH_4_ monooxygenases likewise decreased in warmed soil, albeit not significantly (Fig. 5, Table S7, S19). The metatranscriptomic analyses further revealed a trend of warming-induced decreases in transcripts for NuoF, responsible for NADH oxidation, and a drop in transcription of genes related to aerobic respiration and oxidative phosphorylation (Fig. S3, Table S7).

**Figure 5.**
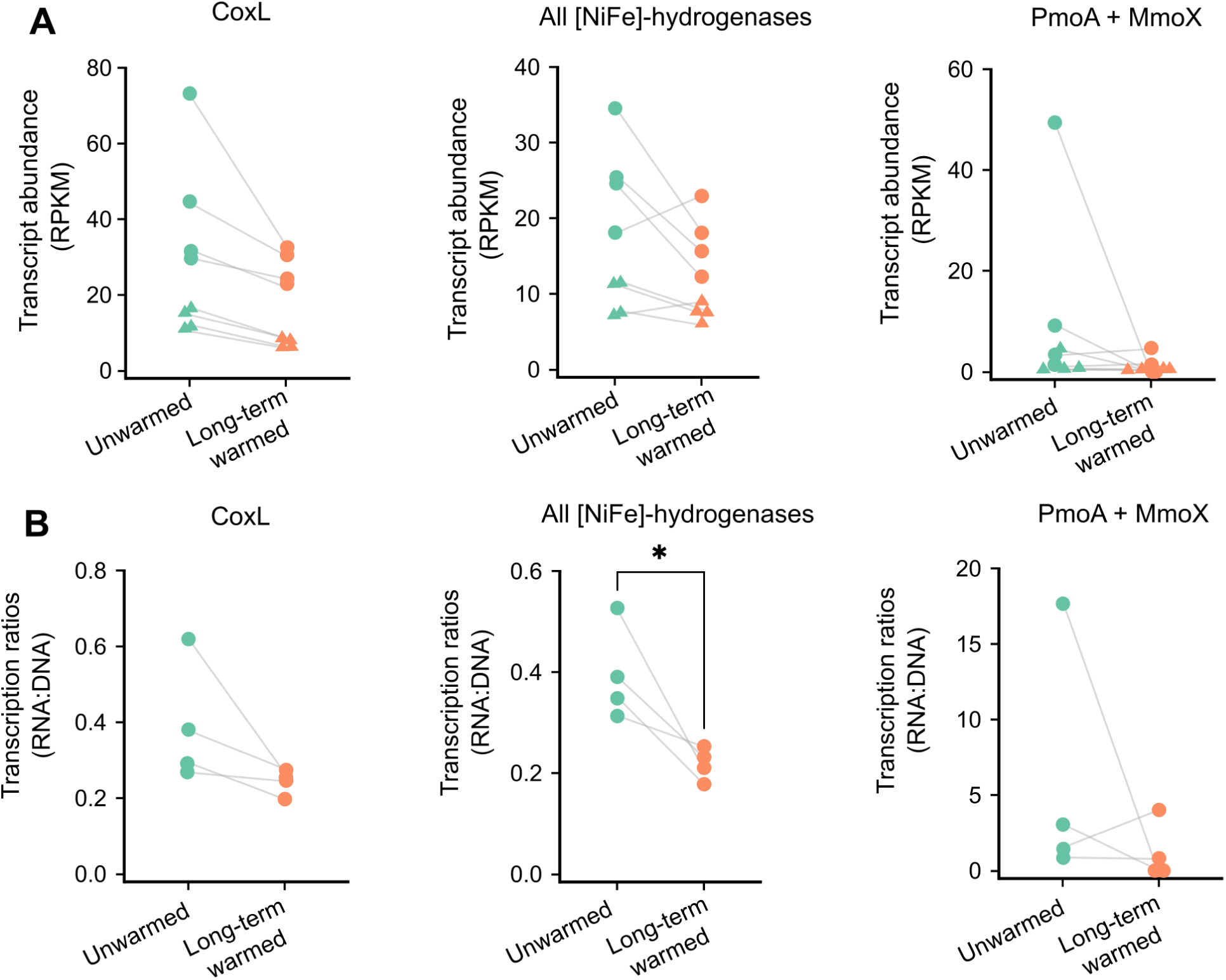
Transcription of trace gas oxidizing genes encoded by the soil community. Transcript abundance of marker genes, calculated as RPKM (**A**), from this study (circles; n= 4) and from public metatranscriptomic short reads (BioProject ID PRJNA663238, triangles; n= 4). Black lines show the mean ± SE (n = 8). Transcription ratio (RNA:DNA) of marker genes based on metagenomic and metatranscriptomic short reads from this study (**B**). Lines indicate pairs within a single warming transect that were used for statistical testing.

To evaluate if the decrease of *in situ* transcription of trace gas oxidation genes can be related to the measured *ex situ* oxidation rates, we converted the latter rates per cell by normalizing the bulk soil consumption rates for each gas by the cell count (as based on qPCR data integrated with metagenome data) and the proportion of respective trace gas oxidizers estimated by metagenomic analysis (Fig. 2D–F, Table S10). The *ex situ* cellular rates closely matched the *in situ* gene transcription, with a general 1.83-, 1.73-, and 3.26-fold decrease in cellular gas uptake for CO, H_2_, and CH_4_, respectively, in long-term warmed soil. Together, our data suggest long-term soil warming decreased trace gas oxidation on a per-cell basis.

### Shifts in relative abundance of trace gas oxidizers are consistent in long-term warmed tundra and temperate forest soils

To determine if our findings are generalisable to other soils experiencing warming, we retrieved and reanalysed metagenomes from two other controlled warming studies, namely tundra soil from the Carbon in Permafrost Experimental Heating Research (CiPEHR) site (warming at 2–5°C above ambient for 1.5 years; AK, USA) (47) and temperate forest soil at the Harvard Forest Long Term Ecological Research (LTER) site (warming at 5°C above ambient for 5, 8, and 20 years; MA, USA) (48) (Table S21). In the tundra soil metagenomes, we detected a 23.1% and 12.6% higher relative abundance of CO and H_2_ oxidation genes per cell, respectively (Fig. 6A, B).

**Figure 6.**
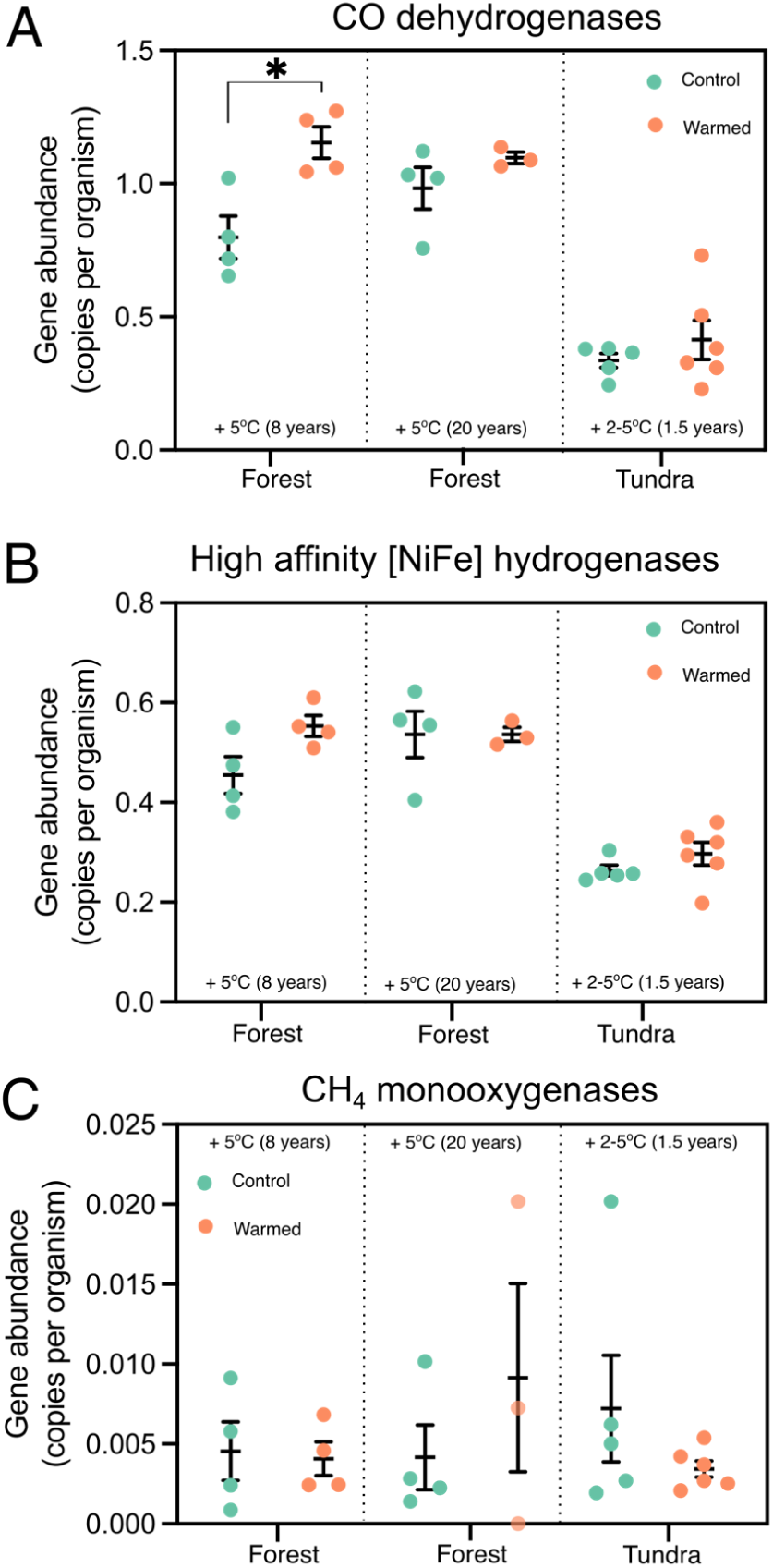
Relative abundance of trace gas oxidizers in global, long-term experimentally warmed soils. Abundances of genes encoding CO dehydrogenases (*coxL*) (**A**),high-affinity [NiFe]-hydrogenases (group 1f, 1h, 1l and 2a)(**B**), and CH_4_ monooxygenases (*pmoA* + *mmoX*) (**C**) in tundra and forest soils under different long-term warming scenarios. The abundance of metabolic marker genes is shown based on metagenomic short reads from two forest soils (48) warmed by +5°C for 8 and 20 years, and a tundra soil (47) warmed by 2–5°C for 1.5 years. The asterisk denotes a significant difference between warmed and control soil (Mann-Whitney test, *p* < 0.05; n= 4). Black lines show the mean± SE.

Additionally, we observed a 2.1-fold decrease in CH_4_ monooxygenase genes under warming (Fig. 6C), which replicates our previous findings. Similar trends were also observed for the LTER temperate forest soil warmed for 5 and 8 years; however, in this soil, methanotroph abundance was 2-fold higher after 20 years of warming (Fig. 6). Together, these results suggest that long-term warming promotes soil H_2_ and CO oxidizers, while it does not seem to consistently promote methanotrophs.

## Discussion

Soil warming has been shown to increase microbial respiration and growth (35), and nitrogen mineralization (49), eventually leading to cumulative carbon and nitrogen losses in topsoils (34, 47, 48, 50). In light of the ability of heterotrophic trace gas oxidizers to supplement energy demand from atmospheric substrates, we initially hypothesized that trace gas oxidizers would increase their gas consumption under carbon- and nitrogen-depleted conditions in long-term warmed soil. Contrary to this hypothesis, our study did not show an increase *in situ* sink capacity for CO, H_2_, and CH_4_ of warmed soils. Moreover, *ex situ* cell-specific oxidation rates of CO, H_2_, and CH_4_ did not increase with long-term (> 60 years) warming of 6°C (Fig. 2D–F). Furthermore, *in situ* community-level transcription of marker genes encoding enzymes for trace gas oxidation showed decreasing trends (Fig. 5). This may be due to the observed non-decreased levels of dissolved organic carbon and nitrogen (Fig. S2), which likely continued to sufficiently support heterotrophy in organisms capable of atmospheric CO and H_2_ oxidation. Although decreased dissolved organic carbon and nitrogen levels have been reported at the same site previously (51, 52), our results indicate that such depletion is not constant, and yearly or possibly seasonal fluctuations occur. We can only speculate that this is connected to seasonal variations in plant activity. The non-decreased levels of dissolved organic carbon and nitrogen are consistent with our *in situ* gas measurements, which show significantly higher respiration rates in warmed soil compared to those in unwarmed soil.

Despite the declining cellular-level responses, the overall *ex situ* sink capacity of warmed soil increased for CO and H_2_. We demonstrated that this effect was possibly driven by the increase of total microbial cell number and biomass in three out of the four studied warming transects, thereby compensating for reduced cellular gas uptake activity under long-term warming (Fig. S2). It should be noted that previous work on the same study site has, in contrast, observed a decrease in microbial biomass and abundance (51), while yet another study shows no change in microbial biomass carbon with warming (52). Thus, the dominant soil CO and H_2_ oxidizing populations appear to retain their function with long-term warming, but multi-season studies across a larger geographical range are needed to definitively confirm this observation. Moreover, the fact that their relative abundance, especially those affiliated with *Actinomycetia*, increased across the three investigated long-term warmed soil ecosystems (Fig. 6) suggests that trace gases likely remain a highly valuable energy source for soil microbes to persist under warmer conditions (23, 53). Given the high capacity for CO and H_2_ uptake across different soil types and the substantial community size of trace gas oxidizers (15), our results suggest that soils can serve as a resilient and reliable sink for these trace gases in a warmer future.

In contrast to CO and H_2_ uptake, the soil sink capacity for the potent greenhouse gas CH4 remained stable, with some indications of weakening due to prolonged warming. This is suggested by multiple lines of evidence, which, although individually not statistically significant, all showed the same pattern of decrease. Firstly, while the aerated soil maintained a net consumption of atmospheric CH_4_, in two out of the four studied warming transects, both *in situ* and *ex situ* rates of CH_4_ uptake declined with long-term warming (Fig. 1B, 2C). Likewise, relative abundance, gene transcription, and DNA:RNA ratios of methanotrophs in two to three out of the four warming transects declined with warming. Finally, in conjunction with the observation of decreased methanotroph abundance in warmed Alaskan tundra soil, all data suggest that the soil CH_4_ sink in upland soils of polar regions is likely to weaken under climate warming. Even medium-term warming of subarctic grassland soil showed a unimodal, temperature-dependent decrease in soil CH_4_ uptake with warming of >4°C (39). Yet, we note that some previous studies on soil warming in Arctic and in alpine grassland soils have reported increased soil CH_4_ uptake (38, 54). In line with these studies, permafrost communities were recently shown to contain relatively more methanotrophs capable of atmospheric CH_4_ oxidation with warming (55). Importantly, previous pure-culture work has demonstrated that responses of methanotrophic bacteria to warming depend on the available methane concentration (56), suggesting that temperature and substrate availability may interact to determine uptake rates. Collectively, this highlights that methanotrophic populations and the soil CH_4_ sink are sensitive and respond variably to warming. Future studies should therefore address potential interactive effects of global warming and atmospheric methane concentrations. Our work highlights the need to profile the crucial CH_4_ sink function of soils in relation to a warmer future climate.

## Materials and Methods

### Experimental sites and soil sampling

The experimental site is a grassland in Grændalur (the “green valley”) near Hveragerði, Iceland (64.0272328°N, 21.1964731°W), which has been naturally warmed for a minimum of 60 years (45) or possibly even before 1708 (57). At this site, geothermal heat within the site’s bedrock has created a natural radiative warming gradient through the soil, which is exploited in permanent study plots along replicated 50 m long transects ranging from unwarmed soil temperatures to +20°C warming at a 10 cm soil depth. In this study, we only focus on the soil temperature range from unwarmed to +6°C. Temperature logger measurements taken in 2013 reveal a constant warming trend following the seasonal temperature cycle at an elevated level (45), as illustrated in Fig. S1 (58). The ecosystem is a grassland dominated by *Agrostis capillaris* on a slightly acidic Silandic Andosol. Soil samples were collected in July 2022 from a depth of 0–10 cm from four unwarmed plots (16°C at time of sampling) and four warmed plots (approximately 22°C at time of sampling), *i.e.*, eight soil samples in total. All samples were homogenized by mixing in sterile sampling bags on site. In parallel, samples for metatranscriptomics were taken (0–10 cm layer) (59), immediately frozen and kept on dry ice until arrival in the lab, where they were stored at -70°C until RNA extraction. Samples for metagenomics, qPCR, and microcosm incubations were cooled until they reached the laboratory, where the former two were frozen at -20°C and the latter kept at 4°C until incubation within one week.

### Microbial biomass, dissolved organic carbon and nitrogen, and pH

Soil pH was measured in a 1:5 (w/v) soil to 10 mM CaCl₂ suspension using a pH meter. suspension using a pH meter. Dissolved organic carbon (DOC) and total dissolved nitrogen (TDN) concentrations were quantified in 1 M KCl extracts (1:7.5 w/v) with a TOC/TN analyzer (TOC-L CPH/CPN, Shimadzu). Microbial biomass C (Cmic) and N (Nmic) were determined using the chloroform fumigation-extraction method (60): Briefly, soils were fumigated with chloroform for 48 h in the dark at room temperature, then extracted with 1 M KCl (1:7.5 w/v) and analyzed as above. Cmic and Nmic were calculated as the difference in extractable C and N between fumigated and unfumigated samples and corrected for incomplete extraction by dividing Cmic by 0.45 and Nmic by 0.54.

### Surface gas fluxes

*In situ* gas fluxes (CO_2_ and CH_4_) at the eight study plots (*i.e.*, four unwarmed temperature and four +6°C warming plots, see above) were measured using a LI-COR gas analyzer with cavity ringdown technique (LI-7810, LI-COR Biotechnology, USA). A PVC chamber (11 cm in diameter × 10 cm in height; 0.95 L total volume) was attached to a collar fixed in the soil to a depth of 2 cm for at least 1 hour in advance, with the remaining aboveground plant parts removed from the plots before the collar placement. The chamber was connected to the gas analyzer. After chamber closure and initial stabilization, the linear concentration increase of CO_2_ and CH_4_ was recorded for 15 minutes at each plot on two separate days to account for day-to-day variation (2022-07-24 and 2022-07-26). Linearity of concentration increase/decrease was monitored in the field. Fluxes of CO_2_ and CH_4_ were calculated from the slope of the linear concentration change in the chamber (R2 > 0.95) as a function of time.

Due to logistical constraints, samples for quantifying *in situ* fluxes of CO and H_2_ were collected from only one of the four warming transects in October 2025 and analyzed in the laboratory using a trace gas analyzer (see Supplementary Materials for a detailed description).

### Soil gas profiles and δ^13^C measurements

Gas samples of 25 mL were taken from the eight study plots with a stainless-steel sampling probe (ø = 3 mm, l = 40 cm) at three soil depths (5, 10, and 20 cm) and injected into pre-evacuated 12 mL vials (Labco Ltd., UK). Two replicates were taken in each depth of every plot, one for analysis of CO_2_ and CH_4_ concentrations and one for analysis of δ^13^C values in CO_2_ and CH_4_. Additionally, an atmospheric gas sample was taken by drawing air above each plot, resulting in a total of 56 samples. Soil temperatures at 5, 10, 15, and 20 cm were measured using a metal probe and a manual thermometer (TM-80N with a K-type thermocouple probe, Tenmars Electronics, Taiwan) during sampling at each plot. Soil moisture was measured in triplicate at all eight plots at a distance of 10 cm using a moisture meter ML-3 Thetaprobe connected to a HH2 moisture sensor (Eijkelkamp, The Netherlands). Gas samples for quantification and δ^13^C measurement of CO_2_ and CH_4_ were treated as follows. Samples were stored at room temperature before analysis of gas concentration by gas chromatography with thermal conductivity (CO_2_) and flame ionization detection (Agilent 7890B) and cavity ring-down spectroscopy (CRDS, Picarro G2201-I with SAM autosampler, Picarro Inc., USA). For CRDS, samples were diluted with synthetic air to <2,000 ppmv CO_2_. The measured isotope ratios were corrected for drift and concentration effects based on the repeated analysis of an in-house standard gas at different dilutions. To correct for concentration effects, we fitted a linear regression between the measured isotope ratio and the inverse concentration, with individual analyses weighted inversely to the standard deviation during measurement to account for the lower precision of measurements from more dilute solutions (61). δ^13^C values of CH_4_ samples with a concentration < 1.2 ppmv after dilution were excluded as reliable measurements are not possible below this concentration. CO, H_2_, and CH_4_ soil concentration values were fitted into a diffusion model to estimate gas concentration across the soil profile (Supplementary Materials). The Rayleigh distillation function, with a fractionation factor (α) of 1.020, was used to predict δ) of 1.020, was used to predict δ^13^C–CH_4_ values.

### Microcosm incubations

Soil microcosms were set up to determine the ability of soil microbial communities to oxidize trace levels of CO, H_2_, and CH_4_, under control, short-term warming, and long-term warming scenarios. Soil samples from four unwarmed plots and four warmed (+6°C) plots were used (sampling and storage described above). To establish the control and short-term warming treatments, approximately 2.5 g of soil from unwarmed plot samples was placed in 160 mL serum vials and incubated at 13°C for the control soil treatment (n= 4) and at 19°C for the short-term warmed treatment (n = 4). Likewise, soil from the warmed plot samples was incubated at 19°C for the long-term warmed treatment (n = 4). Both 13°C and 19°C fall within the range of the respective average summer temperatures in unwarmed plots and warmed plots over the past nine years (Fig. S1). Each serum vial was amended with CO, H_2_, and CH_4_ to initial mixing ratios of ≈ 10 ppmv for each gas. At the start of the experiment, the serum vials were also adjusted to 15% overpressure with ambient laboratory air to compensate for the gas sampled during the experiment. 2 mL of headspace gas were sampled using a gas-tight syringe at different time intervals (eight times in the first three days and eight times in the remaining incubation time) and up to 40 days. Gas samples were measured using a VICI gas chromatograph with a pulsed discharge helium ionization detector calibrated against standard ultra-pure CO, H_2_, and CH_4_ gas mixtures as previously described (22). Pooled autoclaved soil (60 min at 120°C) and empty serum vials were prepared as negative controls.

### Kinetic analysis

For kinetic analysis, measurement time points with individual gas concentrations exceeding 0.4 ppmv were used. The gas consumption patterns were fitted with both linear and exponential models. In all cases, the exponential model provided a better fit to the data based on Akaike information criterion scores; thus, the first-order reaction rate constants were calculated and used for kinetic modeling. The bulk atmospheric gas oxidation rates of each sample were calculated with respect to the mean atmospheric mixing ratio of the corresponding gas (CO: 0.09 ppmv; H_2_: 0.53 ppmv; CH_4_: 1.9 ppmv) (9, 11, 13). Cell-specific gas oxidation rates were determined under the assumption that all cells are viable and active. The soil cell abundance was determined using 16S rRNA gene copy numbers from quantitative PCR (qPCR, see details below) corrected to the average 16S rRNA gene copy numbers derived from the metagenomic short reads (see below). The cell-specific gas oxidation rates were then inferred by dividing the bulk oxidation rate by the product of the estimated relative abundance of trace gas oxidizers derived from the metagenomic short reads (the average gene copy numbers, assuming one copy per cell; see details below) and the soil cell abundance.

### DNA and RNA extraction, qPCR, and sequencing

Community DNA was extracted from ≈ 0.25 g of soil using the DNeasy PowerSoil Pro kit following the manufacturer’s instructions. The number of bacterial and archaeal cells in each sample was estimated using quantitative PCR (qPCR) of the 16S rRNA gene with the universal primer pair F515 and R806, as described previously (62). Further details of processing are described in the Supplementary Materials.

The RNA extraction and metatranscriptomic sequencing were described in (52). Detailed steps are described in the Supplementary Materials.

### Functional annotation of unassembled metagenome and metatranscriptome reads

High-quality DNA short reads and mRNA reads underwent metabolic annotation using DIAMOND v.2.0.15 (63). Reads were aligned against a custom database of 51 metabolic marker proteins (64), covering the major pathways for aerobic and anaerobic respiration, energy conservation from organic and inorganic compounds, carbon fixation, nitrogen fixation, and phototrophy (65). The alignments were filtered to retain sequences of at least 40 amino acids in length or with at least 80% query or subject coverage. They were further filtered based on an identity threshold of 80% (PsaA), 75% (HbsT), 70% (PsbA, IsoA, AtpA, YgfK, and ARO), 60% (CoxL, MmoX, AmoA, NxrA, RbcL, NuoF, [FeFe]-hydrogenases and [NiFe]-hydrogenases group 4), or 50% (all other genes). For metagenomic data, the gene abundance was converted to reads per kilobase million (RPKM) as follows: RPKM = X / total sample reads × 10^6^, where X = reads aligned to a gene / gene length (kbp) (66). This value was divided by the mean abundance of 14 universal single-copy ribosomal marker genes (also in RPKM) provided by SingleM v0.13.2 (67) to obtain the ‘average gene copies per cell’. This value corresponds to the proportion of community members that encode the gene. For metatranscriptomic data, the gene transcription was expressed in RPKM as described above. Heatmaps showing the metagenomic data and RPKM for metatranscriptomes were generated in R version 4.3.1 (68) using the ggplot2 package (69).

### Metagenome assembly and taxonomic analysis

Metagenome reads were processed using Trimmomatic v0.39 (70) to remove adapter contamination, trim low-quality ends of reads, and remove sequences shorter than 36 bp. The quality of the filtered reads was rechecked using Fastp v0.23.4 (71). Filtered reads were assembled individually using MEGAHIT v1.2.9 (72). Gene calling was done using Prodigal v2.6.3 (73) with the mode “-meta”. The same custom database of 51 metabolic marker proteins as mentioned above was used for a DIAMOND blastp search v2.1.9 (*6*3) with metagenome gene calls as queries; best hits with percent identity >75 and query coverage >70 to the CoxL, MmoX, PmoA, [NiFe]-hydrogenases, [FeFe]-hydrogenases, [Fe]-hydrogenases were retrieved. These hits were further compared using DIAMOND blastp v2.0.14 against the bacterial RefSeq database to assign taxonomy to the target gene calls. The taxonomy search utilized filtering for only the best hits, with the following parameters in descending priority: max bitscore > min e-value > max percent identity. The RefSeq r220 database was accessed on 2023-11-20 (*74*) using the NCBI Datasets v15.29.0 tool (http://www.ncbi.nlm.nih.gov/datasets) to retrieve bacterial genomes and MAGs, which were classified by the GTDB taxonomy (bac120_taxonomy_r214). Some of the metagenome gene calls had hits to RefSeq genomes (or MAGs) not included in the GTDB taxonomy; therefore, GTDB-Tk v2.4 (75) was used to assign taxonomy to these genomes. In this way, it was possible to link trace gas oxidation genes assembled in metagenomes with their closest relatives of known taxonomic identity. The relative abundance of target trace gas oxidation genes was expressed as the number of target genes per million total assembled genes in each metagenome. Richness of the closest genomes with respective trace gas oxidation genes was calculated for each sample. Data were processed in R 4.3.1 (68) using the packages tidyverse 2.0.0 (76), tidylog 1.0.2 (*77*), and vegan 2.6-4 (78).

### Metagenomic binning, functional annotation, and phylogenetic analysis

A total of eight metagenomes (4× from unwarmed plots and 4× from warmed plots), which were quality controlled using BBTools (79) as described above and co-assembled using MEGAHIT v1.2.9 (72) with the minimum contig length of 500 bp. Coverage profiles for each contig were generated by mapping the short reads to the assemblies using BBMAP v38.81 (79) and SAMtools v1.19.3 (80). Genome binning was performed with MetaBat2 v2.15.5 (81), MaxBin2 v2.2.7 (82), CONCOCT v1.1.0 (83), and SemiBin2 v2.1.0 (84). The four bin sets were consolidated using the bin refinement module of MetaWrap v1.3.2 (85) and dereplicated using dRep v3.5.0 (86) with a minimum completeness of 50% and contamination of 10%. The completeness, contamination, and heterogeneity of the bins were assessed using CheckM2 v1.0.1 (87), with the quality thresholds selected as previously described (88). Only medium (completeness > 50%, contamination < 10%) and high (completeness > 90%, contamination < 5%) quality bins were retained for further processing and were termed metagenome-assembled genomes (MAGs). Taxonomy was assigned to each MAG using GTDB-Tk v2.4.0 (*7*5) with GTDB release 220 (89). Relative abundance of each MAG in each sample was calculated using the ‘genome’ function (--min-covered-fraction 0, --min-read-aligned-percent 0.75, and –min-read-percent-identity 0.95) of CoverM v0.7.0 (90). Recovered MAGs were then metabolically annotated against the custom database using Prodigal v2.6.3 (73) with the mode “-single” to predict open reading frames (ORFs) and DIAMOND v2.0.15 (63) to identify metabolic marker genes as previously described. A maximum likelihood tree was constructed for MAGs containing trace gas oxidizing genes using GTDB-tk v2.4.0 with the commands ’*identify’* and ’*align’* (75). The tree was constructed using IQ-TREE v2.3.4 (91) with 1,000 ultrafast bootstrap using the LG+F+I+G4 model. The tree was visualized and mid-rooted using the Interactive Tree of Life (iTOL) (92).

### Analysis of publicly available metagenomes from long-term soil warming studies

Given the limited number of samples, we included analyses of public metagenomic (BioProject ID PRJNA746424) and metatranscriptomic (BioProject ID PRJNA663238) data obtained from the same studied site (49). To compare the effects of warming on trace gas oxidation capacity of different types of soils, publicly available metagenomes from temperate forest soil at the Harvard Forest Long Term Ecological Research (LTER) site (MA, USA) (48) and from tundra soil from the Carbon in Permafrost Experimental Heating Research (CiPEHR) site (Alaska, USA) (47) were retrieved. The chosen LTER soil was sampled from 0–20 cm depths, and the warmed soil had been heated consistently 5°C above the temperatures observed in the nearby control soil for 5, 8, and 20 years (48). The soil from the CiPEHR sites was collected from depths of 15–25 cm, with the warmed soil experimentally heated between 2–5°C above the control soil temperature for 1.5 years (47). All datasets were quality-assessed and annotated against 51 metabolic marker proteins, as described above.

### Statistical analysis

All analyses were performed using the statistical software R version 4.3.1 (68) and GraphPad Prism version 10.4.1. Linear mixed models generated with the function lme of the package nlme v.3.1-164 (93) were used to evaluate the influence of short and long-term warming on *in situ* gas fluxes and *ex situ* CO, H_2_, and CH_4_ oxidation rates. The models included the field transect as a random effect to account for the paired sampling design and were checked for meeting the model assumptions by the function check_model of the package performance (94). Oxidation rate values were log-transformed to normalize residuals where necessary. Likewise, linear mixed models were used to test for significant differences in the relative abundance of transcripts and transcription ratios of marker genes for trace gas oxidation, as well as in the relative abundance of MAGs mapped against our metagenomes. Statistical significance was set at ≤ 0.05, and a trend at ≤ 0.1. Detailed statistical evaluation and outputs are provided in Table S11–S12. For public databases, differences in marker gene abundances were determined using the Mann-Whitney test in GraphPad Prism.

## Acknowledgements

We thank Páll Sigurðsson for help during field sampling and Christoph Gall for help with the analysis of soil parameters.

## Funding

VT, JAN, MW, and AD were supported by the JuniorStar project 21-17322M from the Czech Science Foundation to AD.

MBD was funded by the German Science Foundation (DFG, BO 5559/1-1 nr. 433256088).

ATT and AS acknowledge funding from the Research Council of Norway (FRIPRO projects LoAir, 315129, and SHRINK, 344999).

PML acknowledges an Australian Research Council DECRA Fellowship (DE250101210) for salary support.

CG acknowledges an Australian Research Council Future Fellowship (FT240100502) for salary support.

LK acknowledges a Research Council of Finland Fellowship (354501) for salary support.

AR acknowledges funding from the Austrian Science Fund (FWF), grant https://doi.org/10.55776/COE7.

## Author contributions

Conceptualization: AD, CG, VT, DN, PML, MW

Methodology: VT, DN, PML, MW, JAN, KYS, JK, MBD, AS, TS, LK, CB, AR, TU, ATT, CG, BDS, AD

Investigation: DN, VT, MW, JAN, MBD, AS, TS, LK, CG, AD

Visualization: VT, DN Supervision: AT, CG, BDS, AD

Writing—original draft: DN, PML, VT, CG, AD

Writing—review & editing: VT, DN, PML, CB, AR, TU, ATT, CG, BDS, AD, MW, AS

## Competing interests

The authors declare that they have no competing interests.

## Data availability

All data needed to evaluate the conclusions in the paper are present in the paper and/or the Supplementary Materials. The raw sequencing data, together with sample metadata, have been deposited at the ENA under the accession numbers PRJEB98590 (metagenome data) and PRJNA1130687 (metatranscriptome data, Dahl et al., 2025), run accessions SRR29850129, SRR29850130, SRR29850145, SRR29850146, SRR29850149, SRR29850150, SRR29850151, SRR29850152. Processing scripts are deposited at a public repository https://github.com/TlaskalV/trace_gas_oxidation_and_warming/.

## Supplementary Materials

### Supplementary Methods

#### Gas diffusion model

We fitted a diffusion and model assuming steady state conditions (Eq. 1) where *dC_i_/dt* is the concentration change of gas *i* over time in a soil volume [mol m^-3^ sec^-1^], *Di_i_* is the concentration change due to diffusion [mol m^-3^ sec^-1^] and *R_i_* is the concentration change due to microbial production or consumption [mol m^-3^ sec^-1^].

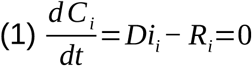

*Di_i_* was calculated following Fick’s second law of diffusion (Eq. 2) adjusted for air-filled porosity (*ε_a_*; dimensionless) and soil tortuosity (*D_S_/D_0_,* i.e., diffusivity in soil relative to air; dimensionless), where *C_i_* stands for the concentration of gas *i* [mol m^-3^], *z* for depth [m] and D_i_ for the diffusion coefficient of gas *i* in air [m^2^ sec^-1^]. In this initial simulation, we assume that soil tortuosity and diffusivity are constant throughout the simulated soil profile.

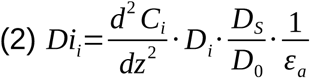

*R_i_* was calculated as a pseudo first-order reaction with a known reaction rate at atmospheric concentration (eq 3), where *R_atm_*is the reaction rate at atmospheric concentration [mol kg^-1^ d.w.], *C_i,atm_* is the concentration of gas *i* in the atmosphere [mol m^-3^], and BD is the bulk density of the soil [kg m^-3^]. This means that we assume that the *potential* soil oxidation rate remains constant with depth, but that the *realized* oxidation rate with changes with depth in response to changing in gas concentration.

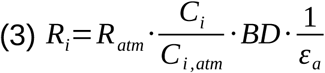

Substituting Eq. 2–3 into Eq. 1 gives

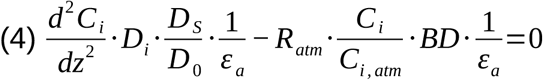

which can be simplified to

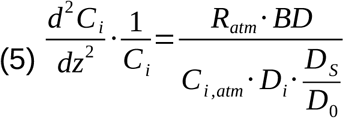

Substituting the right-hand side of Eq. 5 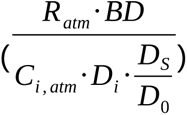 for the constant k gives a second order differential equation

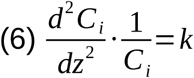

which resolves to

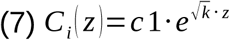

We apply the following constraints, (i) *C_i_(z=0) = C_i,atm_*, i.e., the concentration at the soil surface is the known atmospheric concentration and (ii) *C_i_’(z)<0*, the concentrations decrease with depth. These conditions are only met when *c1=0* and *c2=C_i,atm_*. The depth profile of a trace gas was therefore described as

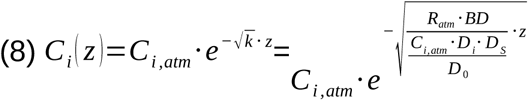

The flux at depth z can be calculated as:

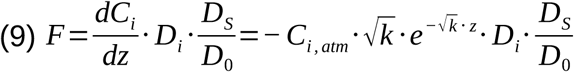

The surface flux (z = 0) is therefore

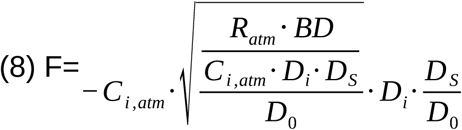

Parameters applied:

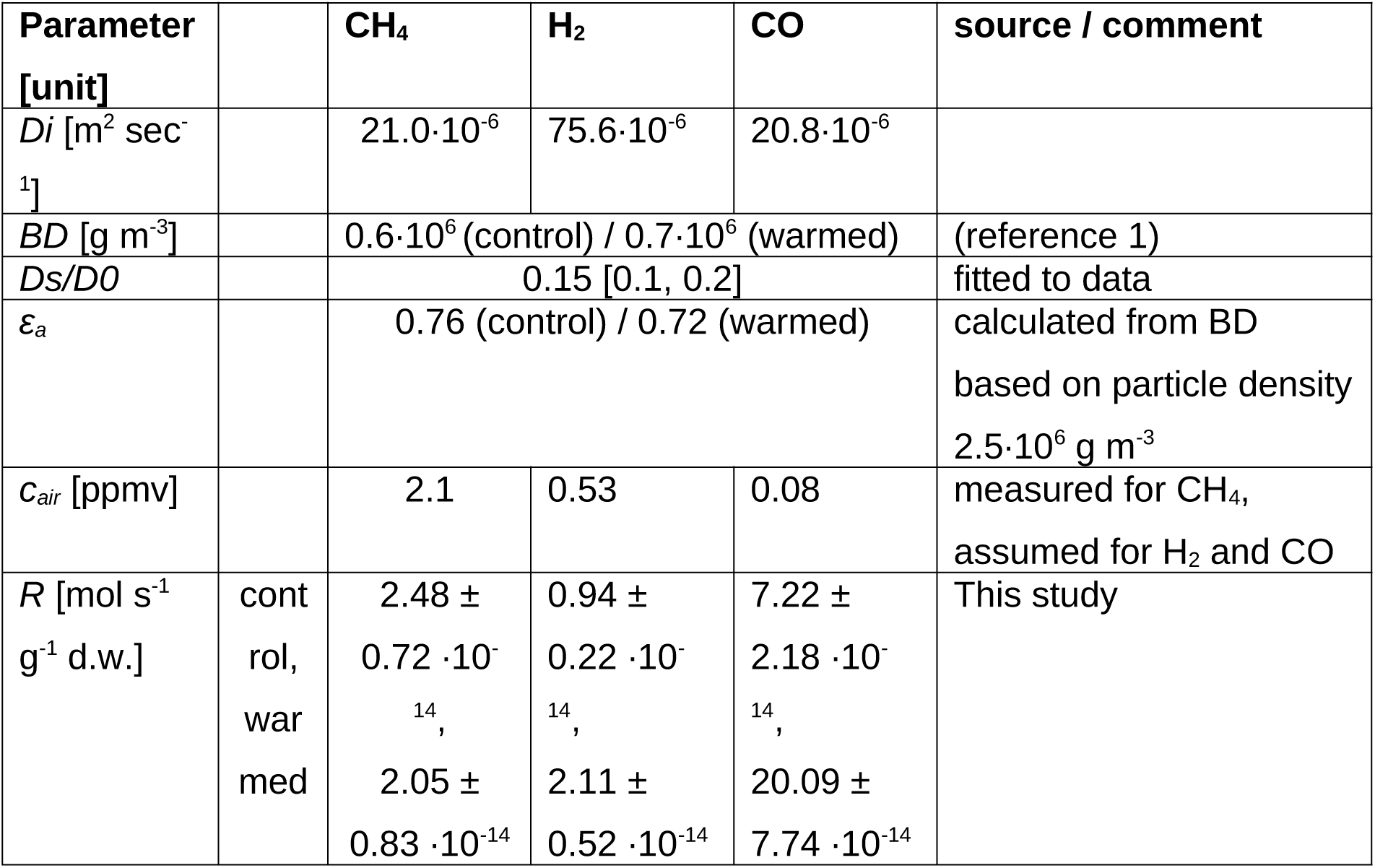

### *In situ* CH_4_, H_2_ and CO uptake

For the *in situ* H_2_ and CO uptake measurements, a metal frame was fixed in the soil of a study plot and the plant material within the frame was removed. Then, an acrylic glass chamber (edge length 20 cm × 20 cm, total volume of 3,600 mL) was placed on the metal frame. Gas samples from the chamber were taken after 0, 2, 4, 8, 16, and 32 min via an embedded butyl rubber stopper using a gas tight syringe (25MDR-LL-GT, SGE International Pty Ltd, Australia). The gas samples were injected into 12 mL vials (738W, Labco, UK) prefilled with ultrapure water, allowing the gas to displace the water. 2 mL of each the gas sample were analyzed using a gas chromatograph (ThermoScientific Trace 1300, Thermo Fisher Scientific) equipped with a sample loop, a Hayesep column (SU12875, RESTEK, Bellefonte, Pennsylvania, USA), a Molsieve 5 A column (PKC17080, RESTEK), a pulsed discharge detector, and a flame ionization detector. A high-quality gas containing 2.5 ppmv CH_4_, 2.5 ppmv H_2_, and 2.5 ppmv CO in N_2_ served as standard (HiQ, AGA, Sweden) (29).

### DNA qPCR, and sequencing

The qPCR assays were performed in triplicate using a 96-well plate in a QuantStudio 7 Flex Real-Time PCR System (Applied Biosystems) and 16S rRNA gene copy numbers were quantified against a serially diluted pMA plasmid containing the Escherichia coli 16S rRNA gene. Metagenomic shotgun libraries, including an extraction blank control, were prepared with the Illumina DNA Prep Tagmentation kit and sequenced on an Illumina NovaSeq 6000 SP at the Ramaciotti Centre for Genomics, University of New South Wales. Two samples (Unwarmed 1 and Warmed 1) were deeply sequenced with an average of 131,415,352 read pairs whereas an average of 35,238,799 read pairs was generated from the other samples. The negative control returned only 139 read pairs, suggesting minimal contaminants. Raw metagenomic data were quality assessed using FastQC v0.11.7 and MultiQC v1.12. The BBDuk function of the BBTools suite v38.96 (79) was used to remove the 151^st^ base, trim adapters, filter PhiX reads, trim the 3’ end at quality threshold of 15 and discard reads below 50 bp in length.

### RNA extraction and sequencing

Soil samples (≈ 40 g) were homogenized (CryoMill, Retsch GmbH, Haan, Germany) under frozen conditions (via liquid nitrogen) and RNA was extracted from each sample (≈ 0.5 g) using the RNeasy PowerSoil Total RNA Kit (Qiagen, Venlo, Netherlands). Extractions were done following the manufacturer’s protocol, but with the following modifications: each soil sample was subjected to two cycles of bead-beating (for lysing and homogenization) with a FastPrep-24 5G (MP Biomedicals, Irvine, CA, USA) to maximize the recovery of RNA per sample. Extracts from both bead-beating steps were combined directly after phase separation in a joint capture tube. Extracts were cleaned with the MEGAclear Transcription Clean-Up Kit (Thermo Fisher Scientific, Waltham, MA, USA). The RNA concentration was determined with the Qubit RNA BR Assay Kit (Thermo Fisher Scientific, Waltham, MA, USA) and the RNA quality was analysed with a Bioanalyzer using the Agilent RNA 6000 Nano Kit (Agilent Technologies, Santa Clara, CA, USA). Sequence libraries were prepared using the NEBNext Ultra II RNA Library Prep Kit for Illumina (New England BioLabs, Ipswich, MA, USA) following the manufacturer’s protocol which included reverse transcription. cDNA input was 100–120 ng. cDNA was fragmented and size selection for 250 bp was performed with HighPrep PCR beads (MagBio Genomics Inc., Gaithersburg, MD, USA). The samples were PE sequenced using a NextSeq 550 System using the NextSeq 500/550 High Output Kit v2.5 (300 Cycles; Illumina, San Diego, CA, USA) at the sequence facility at Greifswald University. Metatranscriptome data were filtered for putative mRNA transcripts using SortMeRNA v.2.1 (95).

---

**Figure S1.**
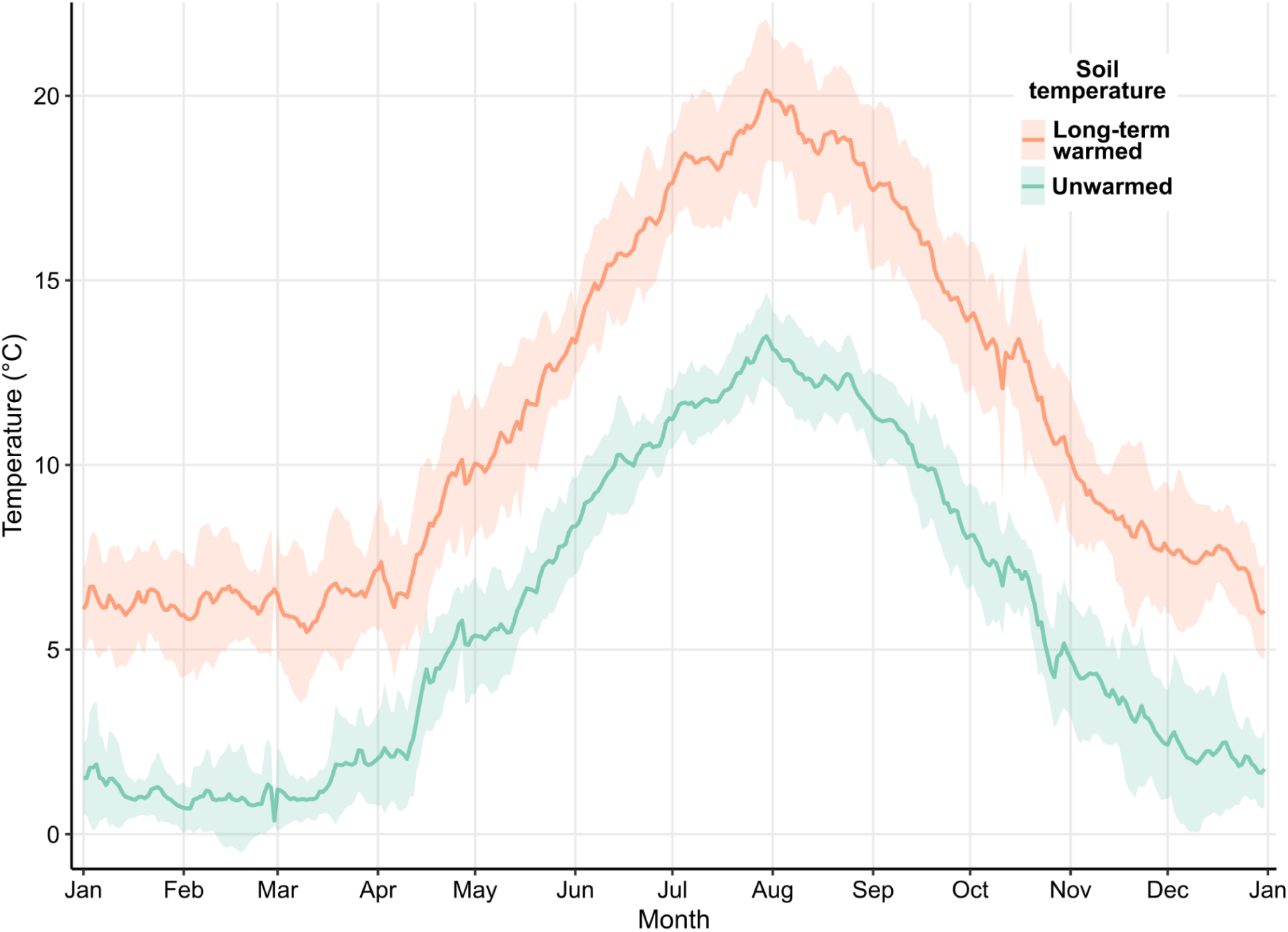
Soil temperatures over the past nine years before sampling. Average annual soil temperature at 10 cm depth for unwarmed (green) and long-term warmed (orange) soil. Lines represent mean temperatures (n = 4), and shaded areas indicate standard deviation across years 2013–2022. Hourly measurements (24.96 ± 0.03 per day, n= 23,837) were first averaged to daily values before calculating annual means.

**Figure S2.**
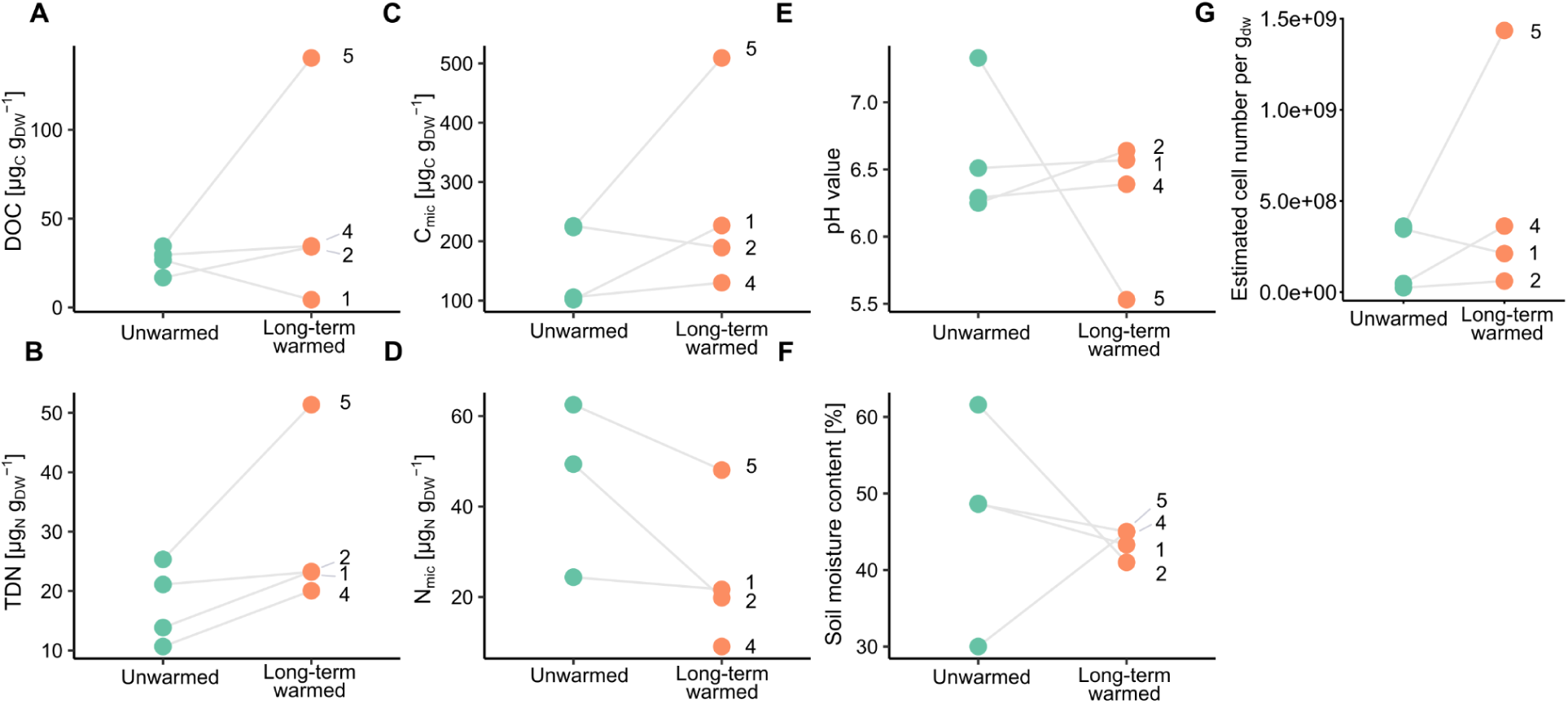
Edaphic characteristics of unwarmed and long-term warmed soil. DOC – dissolved organic carbon (**A**), TDN – total dissolved nitrogen (**B**), C_mic_ – microbial carbon (**C**), N_mic_ – microbial nitrogen (**D**), pH of soil (**E**), field soil moisture content (**F**), cell number per gram of dry soil estimated based on qPCR integrated with metagenome data (**G**). Lines connect pairs within a single warming transect that were used for statistical testing, transect ID is indicated by numbers.

**Figure S3.**
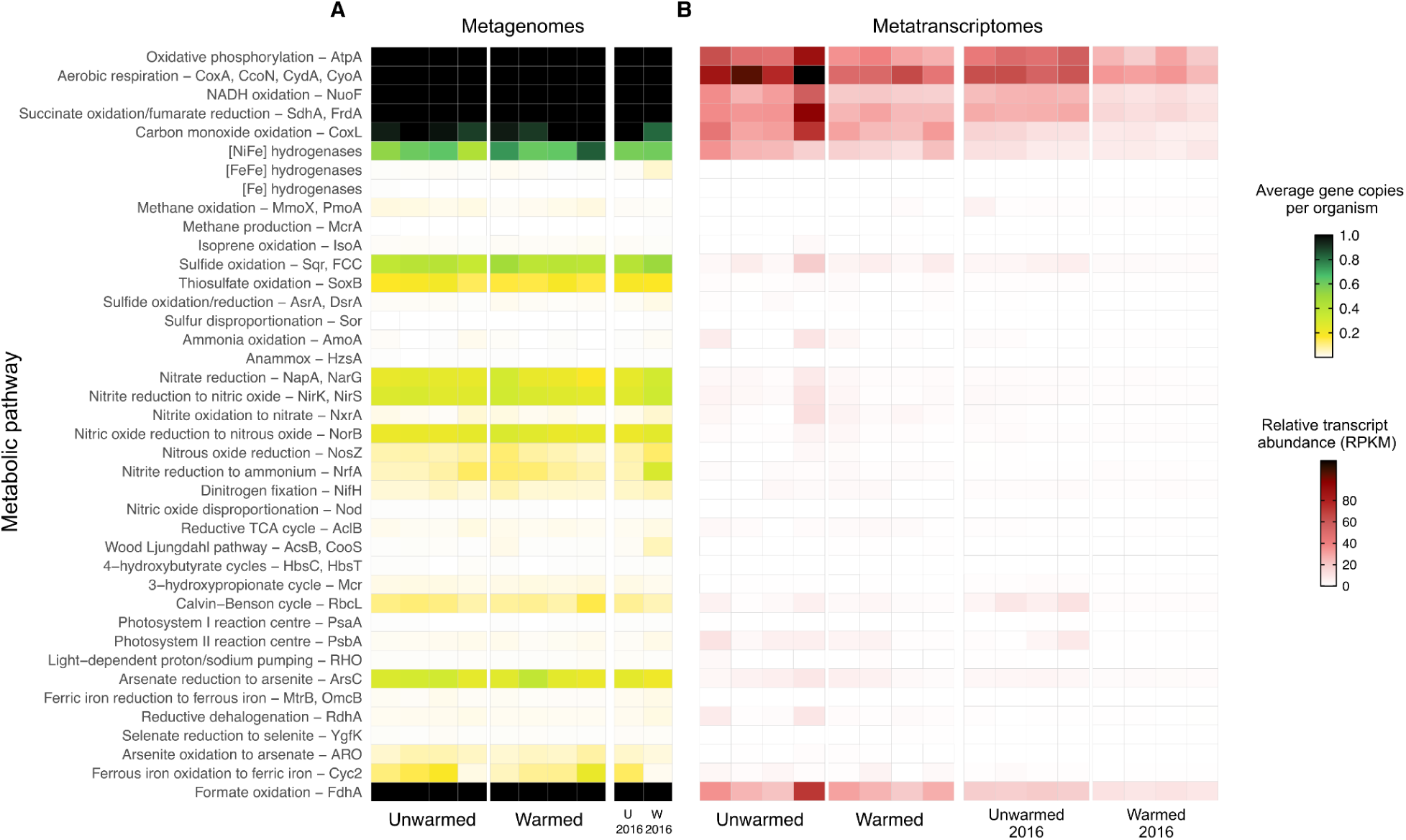
Abundance and transcription of metabolic marker genes encoded by soil community. The abundance of metabolic marker genes is shown based on the metagenomic short reads (A) and metatranscriptomic short reads (B) across unwarmed (n = 4) and warmed plots (n = 4). Publicly available metagenomic data (BioProject PRJNA746424; unwarmed (n = 1) and warmed (n = 1)) and metatranscriptomic data generated in 2016 [BioProject PRJNA663238; unwarmed (n = 4) and warmed (n = 4)] were included.

**Figure S4.**
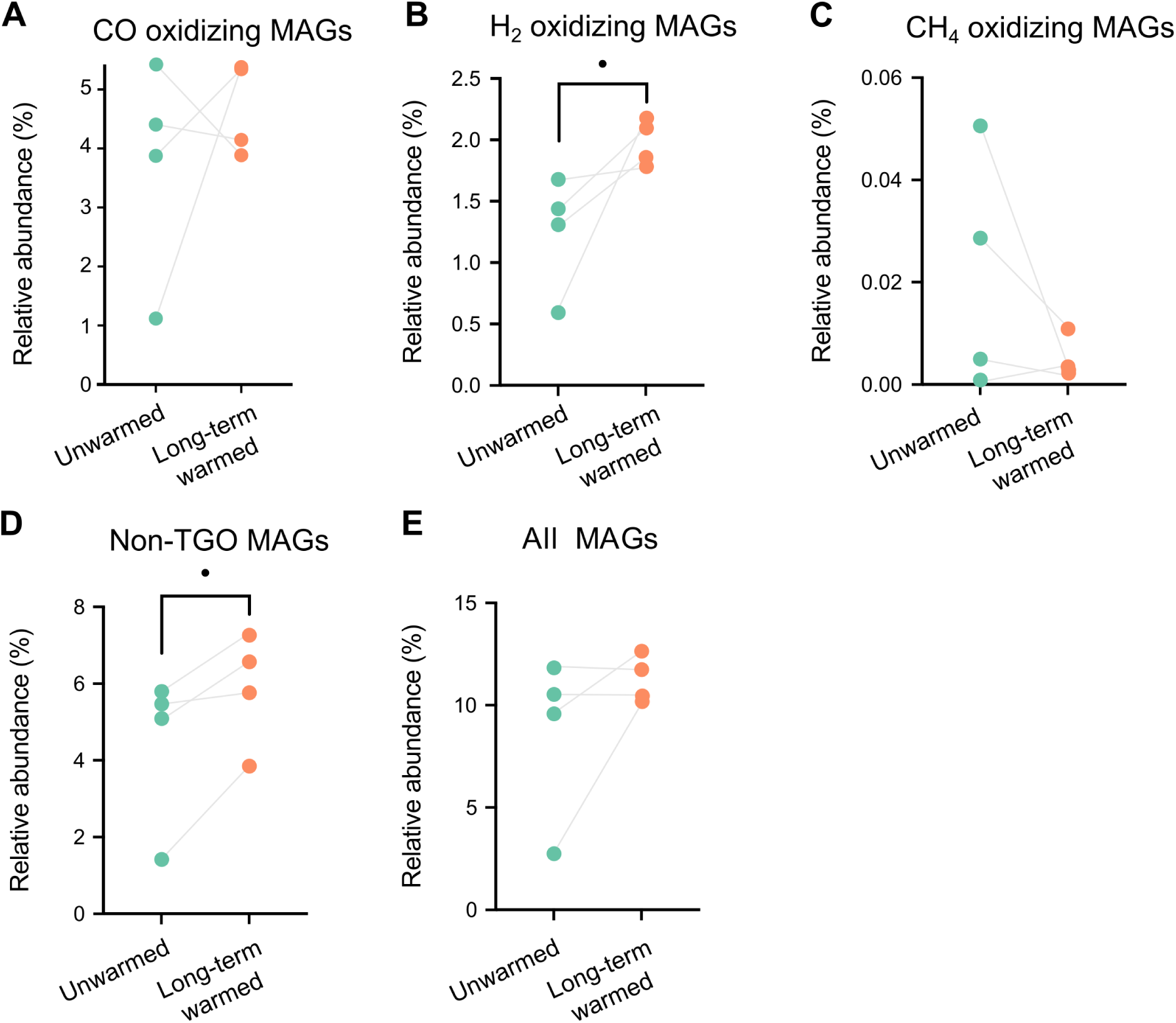
Summed relative abundances of recovered MAGs. Relative abundance of MAGs encoding CoxL (A), high affinity [NiFe] hydrogenases (including groups 1h, 1f, 1l and 2a) (B), and PmoA/MmoX (C) were considered as CO oxidizers, H _2_ oxidizers, and CH_4_ oxidizers, respectively. The rest of MAGs did not encode any of these marker enzymes (D), all MAGs included in the dataset encompass both oxidizers and non-oxidizers (E). Significant differences and trends are depicted by square brackets and asterisk or dot, respectively. Lines connect pairs within a single warming transect that were used for statistical testing.

## Notes

### Competing Interest Statement

The authors have declared no competing interest.

### Summary of Updates

There was an oversight in the author list. This is now corrected.

